# Transcriptomic and DNA methylation modifications during fruit ripening and in response to ABA treatment in sweet cherry

**DOI:** 10.1101/2022.12.02.518926

**Authors:** Nathalie Kuhn, Macarena Arellano, Claudio Ponce, Christian Hodar, Francisco Correa, Salvatore Multari, Stefan Martens, Esther Carrera, José Manuel Donoso, Lee A. Meisel

**Affiliations:** Facultad de Ciencias Agronómicas y de los Alimentos, Pontificia Universidad Católica de Valparaíso, 2340025 Valparaíso, Chile; Universidad de Chile, Instituto de Nutrición y Tecnología de los Alimentos, 820808, Macul, Chile; Instituto de Investigaciones Agropecuarias, Centro Regional INIA Rayentué, 2940000, Rengo, Chile; Fondazione Edmund Mach, Centro di Ricerca e Innovazione, Via E. Mach 1, 38098 San Michele all’Adige, Trentino, Italy; Instituto de Biología Molecular y Celular de Plantas (IBMCP), CSIC-Universidad Politécnica de Valencia, 46022 Valencia, España

**Author notes:** Both authors contributed equally to this work (Nathalie Kuhn, Macarena Arellano). Email addresses. **Highlight** ABA-induced changes in gene expression and methylation in sweet cherry.

**Keywords:** Abscisic acid, bisulfite sequencing, cherries, epigenetic, maturity, non-climacteric, RNA-seq.

## Abstract

Abscisic acid (ABA) plays a key role in the ripening process of non-climacteric fruits, triggering pigment production, fruit softening, and sugar accumulation. Transcriptomic studies show that ABA modifies the expression of several ripening-related genes, but to date, the epigenetic approach has not been utilized to characterize the role of ABA during this process. Therefore, this work aimed to perform transcriptomic and DNA methylation analyses of fruit samples treated with ABA during the fruit ripening process in the non-climacteric sweet cherry model. RNA-seq analyses revealed an overrepresentation of transcripts annotated in functional categories related to ABA response, secondary metabolism, and sugar synthesis. In contrast, Whole Genome Bisulfite Sequencing (WGBS) revealed DNA hypomethylation in the 5’UTR region of genes related to carotene catabolism. Genes encoding xyloglucan enzymes were regulated transcriptionally and epigenetically during ripening. ABA treatment enhanced color development and ripening. GO analysis of DEGs in the RNA-seq of the ABA treatment revealed expression variations in genes encoding members of the Aux/IAA and ARF families. In the WGBS, genes encoding enzymes of the cytokinin biosynthesis had differential DNA methylation after the ABA treatment. Our work shows the genetic factors modulated by ABA at the genetic and epigenetic levels during non-climacteric ripening.

## Introduction

Fruit ripening is an essential process required for seed dispersal and is controlled by a complex network of genetic factors strongly influenced by plant hormones (Karlova et al., 2014; Kumar et al., 2014; Obroucheva et al., 2014). In non-climacteric fruits such as grapevine (*Vitis vinifera L.*), strawberry (*Fragaria × ananassa L.*), and sweet cherry (*Prunus avium*), a characteristic rise in ethylene production and respiratory burst does not occur. Instead, a complex hormone interplay takes place to modulate fruit ripening. The hormones involved in this interplay include abscisic acid (ABA), brassinosteroids (BRs), and auxin, among other phytohormones (Kuhn et al., 2014; McAtee et al., 2013; Fuentes et al., 2019).

The most characteristic feature of non-climacteric fruit ripening is the accumulation of ABA, sugars, and pigments (Wheeler et al., 2009; Luo et al., 2014; Fasoli et al., 2018). Other events occurring during ripening include reducing fruit firmness and decreasing chlorophyll and organic acid content (Ren et al., 2011; Luo et al., 2014; Shen et al., 2014). In this regard, positive biomarkers of non-climacteric ripening are ABA, carotenoids, and anthocyanins, whereas negative biomarkers are associated with carbon fixation, photosynthesis and organic acids (Fasoli et al., 2018).

Ripening involves significant genetic variations (Karlova et al., 2014). In the non-climacteric model sweet cherry, global gene expression analyses in the exocarp of developing fruits have identified several co-regulated genes related to sugar transport and cell wall metabolism (Alkio et al., 2014), which are critical processes of fruit ripening. Anthocyanin accumulation is a characteristic event occurring during the fruit ripening, and yellow and red sweet cherries express several genes of the MYB, bHLH, and WD40 transcription factor families during the anthocyanin biosynthetic process (Wei et al., 2015). Co-expression network analyses in sweet cherry have identified several transcription factors of the bHLH, MYB, WRKY, and ERF families that have similar expression patterns with structural genes of the anthocyanin pathway, such as *PacANS* (Yang et al., 2021). A co-expression analysis conducted at the onset of ripening (*veraison*) of two contrasting grapevine varieties during harvest revealed significant transcriptomic rearrangements that define the reprogramming of berry development (Fasoli et al., 2018). In the same line, a recent work of our group, conducted in two contrasting sweet cherry varieties with differences in the initiation of ripening, showed that they present significant differences at the transcriptomic level, including several genes involved in hormone biosynthesis and signaling, as well as transcriptional regulation (Kuhn et al., 2021b).

Besides global gene expression variation, increasing evidence indicates that epigenetic modifications are crucial during fruit ripening. In this regard, in non-climacteric species, only a few studies explore epigenomic regulation of the ripening process. For instance, in the non-climacteric sweet pepper (*Capsicum annuum* L.), DNA hypomethylation (in the 5’position of cytosine) occurs in the upstream region of the transcriptional start site of several ripening-related genes during fruit ripening (Xiao et al., 2020). In grapevine, it was found that DNA methylation during fruit development represses the expression of genes associated with metabolic and cellular processes, binding, among others (Shangguan et al., 2020). In strawberry fruits, loss of DNA methylation occurs during ripening. Additionally, a DNA methylation inhibitor caused an early ripening phenotype in strawberry, suggesting that DNA hypomethylation is required for fruit ripening (Cheng et al., 2018). This effect differed from the results obtained in oranges, where treatment with a DNA methylation inhibitor delayed ripening, indicating that DNA hypermethylation is required for this process in oranges (Huang et al., 2019).

Despite these works advancing the understanding of non-climacteric ripening, it is still not clear which signals control the genetic and epigenetic changes. In this regard, ABA is an integrator of external and internal cues that regulate gene expression in several plant processes. For instance, in *Arabidopsis*, the MYB96 transcription factor mediates the ABA-auxin signaling pathway under drought conditions (Seo et al., 2009). In the bicolored sweet cherry fruits, hormone-related genes are involved in the anthocyanin regulatory system in response to light, including ABA signaling genes (Guo et al., 2018). In sweet cherry, the expression of several *PavBBX* genes, encoding transcription factors with a similar pattern to the anthocyanin biosynthesis genes, are strongly induced by ABA and other hormones (Wang et al., 2021).

Regarding fruit ripening, a high-throughput microarray analysis of the strawberry fruit development revealed that ABA regulates the expression of most ripening-related genes (Medina-Puche et al., 2016). However, evidence of ABA involvement in epigenetic control during fruit ripening is scarce. Recent work in strawberry fruits showed significant changes in the methylation of adenines in the CDS of mRNAs at ripening initiation, including those transcripts coding for ABA biosynthetic enzymes and signaling factors (Zhou et al., 2021). Considering all these works, ABA could be fundamental in controlling ripening through genetic and epigenetic rewiring.

ABA plays a central role as a positive regulator of ripening in non-climacteric fruits (Fenn and Giovannoni, 2021; Jia et al., 2011; Luo et al., 2014; Setha et al., 2012; Sun et al., 2010; Wheeler et al., 2009). Using a molecular approach in sweet cherry fruits, Shen and coworkers (2014) showed that RNAi lines with reduced expression of *PavNCED1* (a putative 9-*cis*-epoxy-carotenoid dioxygenase gene important for ABA production) have reduced anthocyanin concentrations. Supporting this finding, fruits from the *PavCYP707A2* interference lines, having increased ABA levels, accumulate more anthocyanins than the control lines (Li et al., 2015). In strawberry, virus-induced gene silencing of a putative ABA receptor gene produces an uncolored phenotype that was not reversed by ABA (Jia et al., 2011). In addition, ABA also affects genes related to the cell wall (Karppinen et al., 2018; Zhang et al., 2021).

Maturity time depends on the timing of ripening initiation. In sweet cherry, the difference in harvest time between varieties occurs at the last growth phase – i.e., from the onset of ripening to maturity (Gibeaut et al., 2017). In grapes, maturity advancement depends on the change in fruit color (*veraison*) rather than the increase in the rate of ripening (Cameron et al., 2021). In this regard, ABA can promote several physiological processes in non-climacteric species such as grapevine (Wheeler et al., 2009) and sweet cherry (Kuhn et al., 2021a). However, how ABA controls the timing of ripening initiation at the transcriptomic and epigenetic levels in non-climacteric fruits is still elusive.

Fruit tree models may help improve the understanding of ripening. In this regard, the sweet cherry has risen as a non-climacteric fruit tree model since it has a limited ripening window (Gibeaut et al., 2017), and there are varieties with contrasting maturity times (Kuhn et al., 2021b). Considering that whole-genome sequencing (Shirasawa et al., 2017) and re-sequencing of *Prunus avium were* performed recently (Xanthopoulou et al., 2020, Sharpe et al., 2022), the genomic data in this species is highly accurate. In sweet cherry, though, a small respiratory peak occurs from the de-greening to the pink color initiation stage (Zhao et al., 2013). Variations in compounds that characterize sweet cherry fruits are the decrease in some phenolics and flavonols during development; and the increase in the anthocyanin concentration at the pink or red stages (Teribia et al., 2016; Ponce et al., 2021). This increase occurs simultaneously or soon after ABA accumulation (Ponce et al., 2021). However, there is a cultivar-dependent increase in IAA and GA_4_ (Ponce et al., 2021).

Some important ABA biosynthesis and signaling genes, such as *PavNCED1*, *PacCYP707As*, *PacPYLs*, *PacSnRK2s* and *PacPP2Cs,* are differentially expressed during sweet cherry fruit ripening (Li et al., 2015; Wang et al.,2015), and changes in ABA signaling-related genes precede the upregulation of structural genes of the anthocyanin pathway (Kuhn et al., 2021a). *Ex planta* treatments with ABA promoted fruit coloration and changed ripening-related parameters such as fruit size, soluble solids content, acidity and firmness (Ren et al., 2010; Ren et al., 2011; Shen et al., 2014; Shen et al., 2017; Wang et al., 2015). ABA treatments *in planta* under field conditions at the color initiation stage showed an advance in IAD (a maturity index) and more anthocyanin content at harvest, which was accompanied by the upregulation of genes of the ABA pathway and the anthocyanin route (Kuhn et al., 2021a).

Considering that sweet cherry is a promising model for unraveling the main features of fruit ripening, this work aims to determine the effect that ABA applied *in planta* at ripening initiation has at the genetic and epigenetic levels. For this, we analyzed the effect of ABA on the physiological features of the ripening process in sweet cherry fruits, such as essential marker variations, including hormones and metabolites. Then we analyzed the ripening process at the genetic and epigenetic level by identifying differentially expressed genes (DEGs), differentially methylated genes (DMGs), and overrepresented categories through Gene Ontology (GO) analyses. Finally, we identified GO categories overrepresented in the ABA treatment in the transcriptomic and DNA methylation analyses. This work sheds light on the genetic and epigenetic mechanisms that regulate the ripening process driven by ABA in a non-climacteric species.

## Materials and Methods

### Plant material

Pitted fruits from adult trees of *Prunus avium* L., cultivar ‘Lapins’, grown in a commercial orchard in Rengo, Chile, Lon: O70°43’6.78” Lat: S34°27’16.92”, were used for these studies. All trees underwent standard agronomic practices and had healthy tree vigor. Field measurements and experimental activities were conducted during the 2017-2018 season. Full flowering (50% open flowers) was set as 0 DAFB (days after full bloom). RNA sequencing (RNA-seq) was made on fruits from the Early Pink stage (56 DAFB; Nov 16, 2017), Late Pink stage (60 DAFB; Nov 20, 2017), and Red stage (67 DAFB; Nov 16, 2017) of sweet cherry fruits. Whole Genome Bisulfite sequencing (WGBS) was performed on fruits from the Early Pink and Red stages. Phenolics, hormones, and pigments were quantified at the Red stage. Color and ripening parameters were measured at harvest (69 DAFB; Nov 29, 2017).

### ABA treatment

Four control trees and four ABA-treated trees were chosen randomly in the field. For the treatment, 400 mg L^-1^ of ABA solution (commercial product ProTone®, SL - Sumitomo Chemical Chile) was sprayed until runoff to the tree canopy in the late morning. As previously reported, ABA was applied when fruits transitioned from the straw-yellow to the pink stage (Luo et al., 2014; Time et al., 2021). The day of the treatment was defined as the time point zero (T0, 56 DAFB); control trees were untreated.

RNA-seq analyses were performed comparing ABA-treated fruit collected minutes before the ABA treatment (ABAT0), two hours post-ABA-treatment (ABAT2h), four days post-ABA-treatment (ABAT4), and 11 days post-ABA-treatment (ABAT11) to control fruit samples at similar time points (CT0, CT2h, CT4, CT11, respectively). WGBS analyses were performed on samples from ABA-treated fruits minutes before the treatment (T0) and 11 days post-ABA-treatment (T11) in control and ABA-treated fruits. Fruit samples of ABA-treated and control trees (each sample as a pool of eight fruits per tree) were harvested, frozen immediately in liquid nitrogen, and stored at -80°C. Before metabolite analyses, samples were lyophilized in a LABCONCO® Freeze dry system.

### Physiological evaluations

Physiological assessments at harvest were performed by collecting fruits at 69 DAFB. Color distribution using the CTIFL scale, IAD (Index of Absorbance Difference), soluble solids content, acidity, firmness, and weight were quantified (Chavoshi et al. 2014). For each tree (n = four), 25 fruits were randomly collected. For color distribution, CTIFL sweet cherry color chart (CTIFL, France) was used to allocate the 25 fruits into color categories (category “1” being the lightest and “4” the darkest). VIS/NIF Cherry Meter device (T.R. ® Turoni, Italy) was used for IAD. For width, the equatorial diameter (from the suture of the fruit) was measured with a caliper. For weight, a portable mini scale was used. Firmness was determined by using a durometer (Durofel T.R. ® Turony, Italy) pressed on both cheeks of the fruits and averaging the value (ranging from one to 100%). For soluble solid content and acidity, the five most homogenous fruits (in color and size) were selected and measured using a PAL-BX|ACID Pocket Sugar and Acidity Meter (ATAGO USA, Inc.), which measures degrees Brix and acidity as total malic acid content, respectively.

### Chemicals

Deionized water was purified with the Arium® purification system (Sartorius AG, Goettingen, Germany). Methanol was of LC-MS grade. Authentic standards of phenolic compounds were purchased from Sigma-Aldrich (Milan, Italy), TransMIT PlantMetaChem (Gießen, Germany), and Phytolab GmbH & Co (Vestenbergsgreuth, Germany). The 0.22 μm PTFE membranes were purchased from Merck Millipore, Darmstadt, Germany.

Analytical-grade laboratory reagents were purchased from VWR (Barcelona, Spain) for hormone quantification. Methanol, acetic acid, and acetonitrile were of HPLC LC-MS grade. Authentic standards of ^2^H_6_-ABA, ^2^H_6_-GA1, ^2^H_6_-GA4, and ^2^H_6_-IAA were purchased from commercial house Olchelm (Olomouc, Czech Republic).

### Phenolic compounds quantifications by Ultra High-Performance Liquid Chromatography coupled to Mass Spectrometry (UHPLC-ESI-MS/MS)

For phenolic compounds analyses, around 150 mg of each lyophilized sweet cherry fruits sample was transferred into 15 mL falcon tubes, and a volume of 4 mL of 80% methanol was added to each sample. The samples were sonicated for 20 min, mixed by orbital shaking for three hours at room temperature, and kept overnight at 4°C in the dark. Then, samples were centrifuged (10 min; 1800×g; 4 °C), filtered through 0.220μm PTFE membranes, and stored at − 80 °C until MS analysis. Three biological replicates were used for each sample (T4 -Late pink- and T11 -Red-stages).

Targeted UPLC was performed on a Waters Acquity system (Milford, MA, USA) consisting of a binary pump, an online vacuum degasser, an autosampler, and a column compartment. Separation of the phenolic compounds was achieved on a Waters Acquity HSS T3 column 1.8 μm, 100 mm × 2.1 mm, kept at 40°C. The analysis of phenolic compounds was performed according to the method described by Vhrovsek et al. (2012). Anthocyanins were analyzed using the method described by Arapitsas et al. (2012). A Waters Xevo TQMS instrument equipped with an electrospray (ESI) source was used for Mass spectrometry detection. Data processing was performed using the Mass Lynx Target Lynx Application Manager (Waters).

### Quantification of hormones by UPLC-ESI-MS/MS

For hormone quantification, around 10 mg of dried sweet cherry fruit samples were suspended in 80% methanol-1% acetic acid solution containing a mix with the internal standards, ^2^H_6_-ABA, ^2^H_6_-GA_1_, ^2^H_6_-GA_4_, ^2^H_6_-IAA and the samples were shaken for one hour at 4°C and then maintained at -20°C overnight. The samples were centrifuged, and the supernatant obtained was dried in a vacuum evaporator to be dissolved in 1% acetic acid. A reverse phase column (Oasis HLB) was used to pass the solution as described by Seo et al. (Seo et al., 2011). The eluate was dried and dissolved in 5% acetonitrile-1% acetic acid. An autosampler and reverse phase UPLC chromatography column (2.6 µ m Accucore RP-MS, 100 mm length x 2.1 mm i.d.; ThermoFisher Scientific) was used to separate ABA with an acetonitrile (2%-55%) gradient containing 0.05% acetic acid, at a rate of 400 µ L/min over 22 minutes. Q-Exactive mass spectrometer (Orbitrap detector; ThermoFisher Scientific) was used for ABA detection in a targeted Selected Ion Monitoring (SIM) mode. The ABA concentrations in the samples were obtained with calibration curves and the Xcalibur 4.0 and TraceFinder 4.1 SP1 software programs.

### Carotenoids and chlorophylls quantification by spectrophotometry

For carotenoid and chlorophyll pigment extraction, the method described by Moreno et al. (2013) with modifications was used. Briefly, between 50-100 mg of lyophilized tissue of sweet cherry samples were used. The tissue was ground in liquid nitrogen, and 1.5 mL of absolute methanol was added. The samples were shaken in an orbital shaker for at least two hours in 2 mL Eppendorf tubes, protecting the samples from the light. The samples were centrifuged at 7,840 x *g* for 5 min, and the supernatant was used for the quantification by spectrophotometry using Infinite ® 200 NanoQuant. The quantification was calculated using equations Lichtenthaler and Buschmann (2001) described for absolute methanol as solvent.

### RNA extraction and RNA sequencing

For RNA-seq analysis, samples were collected as described in the “ABA treatment” section. RNA was extracted from 24 fruit samples, with three replicates for each time point: CT0, ABAT0, C2h, ABAT2h, CT4, ABAT4, CT11, and ABAT11.

Total RNA was isolated from 1 g of ground, pitted fruit samples (mesocarp and exocarp-enriched tissue) using the CTAB method with minor modifications, according to Meisel et al. 2005. GeneJET RNA Cleanup and Concentration Micro Kit (ThermoScientific, San Diego, CA, USA) were used for purifying the RNA samples. Purity values (A260/230 and A260/A280) ranged between 1.8–2.2 in all the samples.

For RNA-seq analysis of the fruit samples, one µ g of RNA with RIN (RNA Integrity Number) > 7.0 was used to generate cDNA libraries of the fruit samples using the Illumina TruSeq Stranded mRNA LT Sample Prep Kit, according to the manufacturer’s instructions. The libraries were sequenced in an Illumina platform (Illumina NovaSeq6000 at Macrogen Inc.), and 100 bp paired-end reads were generated. Data was generated using the base-calling Software RTA (Real-Time Analysis) and the Illumina package bcl2fastq.

### Analysis of differentially expressed genes

The data was high quality (the bases with a Q score > 20 represented more than 97% of the reads in all the samples) and free of adaptors. Sequence mapping to the sweet cherry reference genome *Prunus Avium* v1.0.a1 (Shirasawa et al. 2017, available on NCBI) was performed using STAR software (Dobin et al. 2012), setting the parameters by default. Transcript abundance was fitted using a general linear model (GLM), and differential expression between treatments was tested with the limma package (Ritchie et al., 2015). Briefly, the model’s coefficient for each gene represented the estimated fold change between control and treated samples. Moderated t-statistics were computed using these values, interpreted as an ordinary t-statistic except for the standard errors moderated across genes using a simple Bayesian model. Raw *p*-values were corrected for multiple testing using the Benjamini–Hochberg procedure. Differentially expressed genes (DEGs) had an absolute fold-change value of at least two between comparisons, with a *p*-adjusted-value < 0.05. A PCA analysis was performed to evaluate the biological replicates (data not shown).

Functional annotation of *P. avium* reference transcripts (Shirasawa et al. 2017) was obtained from the NCBI database. The Gene Ontology (GO) annotation of the *P. avium* transcripts was performed primarily by a tBLASTx from the *P. avium* mRNA database (NCBI) against the protein sequences from all Rosaceae species available on the Universal Protein Resource (UniProt). The best hit was determined using the following parameters: Percentage identity > 60%, e-value < 1e-20, and the sequence with the highest score. The GO annotations associated with the protein from the Rosaceae species were assigned to the homolog mRNA of *P. avium* using code programming in python.

The GO enrichment analysis was performed with the topGO package in R (Alexa and Rahnenfuhrer, 2021). The GO overrepresented categories were obtained with the Fisher’s Exact Test Enrichment Analysis considering a *p*-value < 0.05. The *p*-value was adjusted using a false discovery rate (FDR) with at least a 95% significance. Furthermore, to reduce redundancy and to group similar GO categories, the platform ReViGO was used (Supek et al. 2011), with the following parameters: a resulting small list (0.5), associated *p-*values from GO enrichment analysis, whole UniProt species as database, and Resnik algorithm for semantic similarity.

### DNA extraction and Whole-Genome Bisulfite Sequencing (WGBS)

CT0 and CT11 sets were used to evaluate methylation variation during fruit ripening, whereas CT11 and ABAT11 were used to assess methylation variation in response to the ABA treatment at T11. Three sample sets were used for methylation analysis (described in the “ABA treatment” section), with three trees as biological replicates.

Genomic DNA was isolated from 0.5 g of ground mesocarp and exocarp-enriched tissue from fruit samples using a CTAB-based method (Healey et al., 2014; Inglis et al., 2018; Aboul-Maaty and Oraby, 2019). Briefly, the samples were extracted with 7 mL of the CTAB 3% buffer by adding 1% β-mercaptoethanol and 1% PVP. After incubation, an equal volume of chloroform:isoamyl alcohol (24:1 v/v, Chl-AIA) was added, mixed, and the aqueous phase was extracted by centrifugation at 8,000x*g* for 15 min. The aqueous phase was incubated with 5 uL of RNAse A at 37 °C for 15 minutes. After, an extraction with Chl-AIA was performed. 1/10 volume of sodium acetate 3M (pH 5.2) and 2/3 volume of cold isopropanol were added and mixed by inversion. The samples were incubated at -20°C for one hour and then centrifugated at 8,000x*g* for 15 min. The aqueous phase was removed by inverting the tubes, and the pellet was cleaned with 1 mL of cold ethanol 70%. The samples were centrifugated at 8,000x*g* for 5 minutes, and the ethanol was removed. Finally, the precipitated DNA was suspended in 100 µ L of molecular-grade water. Using an Infinite M200 PRO Nano Quant (TECAN), spectrophotometric analysis was used to determine DNA concentrations. The integrity of the DNA was visualized on an 0.8% agarose gel.

Approximately one µ g of Genomic DNA from each sample was used to perform Whole Genome Bisulfite Sequencing. Bisulfite libraries were prepared using the EZ DNA Methylation Gold kit (Zymo, CA, USA) and Accel-NGS Methyl-Seq DNA Library Kit (Swift Biosciences, USA). The bisulfite libraries were sequenced on Illumina HiSeq 150bp paired-end.

### Analysis of differential methylation regions and differential methylation genes

For data analysis, paired-end sequencing reads were filtered with TrimGalore to remove Illumina adapters, low-quality bases (quality score < 20), and clonal reads. Filtered reads were mapped to sweet cherry reference genome *Prunus Avium* v1.0.a1 (Shirasawa et al. 2017, available on NCBI) using the bisulfite mapping tool Bismark (Krueger and Andrews, 2011, available on www.bioinformatics.bbsrc.ac.uk/projects/bismark/). The methylation percentage of three methylation contexts (CpG, CHG, and CHH, where H= A, T, or C) were extracted from the bismark alignments using the methylKit R package (Akalin et al. 2012, URL http://code.google.com/p/methylkit/). A PCA analysis was performed to evaluate the correlation between biological replicates (data not shown) Differentially methylated regions (DMRs) were identified from cytosines with a coverage of at least five reads in all the samples through a 200bp sliding-window approach and in intervals of 50 bp (with at least ten methylated cytosines per window). Logistic regression was performed to determine the differential methylation between comparison windows of CT0/CT11 and CT11/ABAT11. The correction of *p*-values to *q*-values was through the sliding linear model (SLIM) method. Windows with a percentage of differential methylation higher than 15% and a *q*-value < 0.05 were identified as differentially methylated regions. Genes with a DMR located in a gene or within 2000 bp upstream of the gene were defined as differentially methylated genes (DMGs). The GO enrichment analysis was performed as described above for RNA-seq analysis.

### In silico analysis of the DEGs/DMGs promoter regions

The gene promoter sequences 2,000 bp upstream (DEGs and DMGs) were obtained from the *P. avium* genome available in the NCBI database using code programming in python. When the promoter region was not available or incomplete, a tBLASTn against the *P. avium* genome cv. Tieton (Wang et al. 2020) was performed in the Genome Database for Rosaceae -GDR, http://rosaceae.org-(considering the higher score sequence, percentage identity > 70%, and e-value < 1e-20), and the sequence was retrieved from GDR by using the JBrowse tool. Hormone-responsive *cis*-elements were predicted by using the PlantCARE database (Lescot et al. 2002, http://bioinformatics.psb.ugent.be/webtools/plantcare/html/).

### Data visualization and statistical analysis

Physiological data parameters were analyzed using the Software GraphPad Prism v6.0. The statistical differences for each parameter were determined with Fisher’s protected least significant difference test (LSD) between ABA treatment and control treatment, using the InfoStat Software (https://www.infostat.com.ar/index.php).

The data for the accumulation of phenolic compounds, hormones, and pigmented compounds were represented as the mean ABA treatment value divided by the mean control treatment value for each compound, and the logarithm in base two was calculated. The bar plots were performed with those values in R 4.0.2 (R Core Team, 2020). The statistical differences were determined with Fisher’s protected least significant difference test (LSD) between ABA treatment and control treatment, using the InfoStat Software (https://www.infostat.com.ar/index.php).

GO overrepresented categories, and the DEGs or DMGs among these GOs were grouped with at least one other category after ReViGO (Supek et al., 2011) analysis and visualized using treemap package in R (v2.4-3, Tennekes M., 2021). Then, the DEGs or DMGs annotated in those categories were visualized in a heatmap using the ggplot2 package in R (Wickham H., 2016). Venn diagrams were designed in R4.0.2 (R Core Team, 2020).

Representations of the promoter regions were designed using exon-intron GraphicMaker (Bhatla N., 2012, available at http://wormweb.org/exonintron) and modified using Gimp software (v2.10, GIMP Development Team, 2019).

## Results

### ABA modulation of ripening-related parameters

In the midseason maturing sweet cherry cultivar’ Lapins’, the modulation of ripening-related parameters in response to ABA treatment was analyzed at the Early Pink and Red stages of fruit development. At harvest, ABA-treated trees produced fruits with increased color homogeneity and intensity (Fig. 1A), reducing CTIFL dispersion values (Fig. 1B) and increasing IAD levels (Fig. 1C), respectively. These “ABA-treated fruits” were also significantly smaller (reduced width and weight), less firm and had a lower percentage of soluble solids content (Supplementary Fig. S1 at JXB online). Several metabolites are markers of the sweet cherry fruit ripening process, including carotenoids, ABA, flavonoids, and anthocyanins (Ponce et al., 2021). Comparative analyses of pigments and hormones at the Red stage of fruit ripening revealed several significant changes in these metabolic ripening markers in fruits from ABA-treated trees compared to untreated. The levels of taxifolin (a flavonoid), total anthocyanins, and total carotenoids were significantly more abundant in the “ABA-treated fruits” compared to untreated (Fig. 2A and B). Additionally, three anthocyanidin glycosides (cyanidin-3-O-galactoside, cyanidin-3-O-sambubioside and peonidin-3-O-glucoside) and “Total other flavonoids” were significantly more abundant in the “ABA-treated fruits” compared to untreated. ABA treatment also reduced chlorophyll a and increased chlorophyll b in the fruits, though not significantly (Fig. 2B). Furthermore, ABA treatment also significantly increased the endogenous IAA and GA_4_ levels in fruits (Fig. 2C).

**Figure 1.**
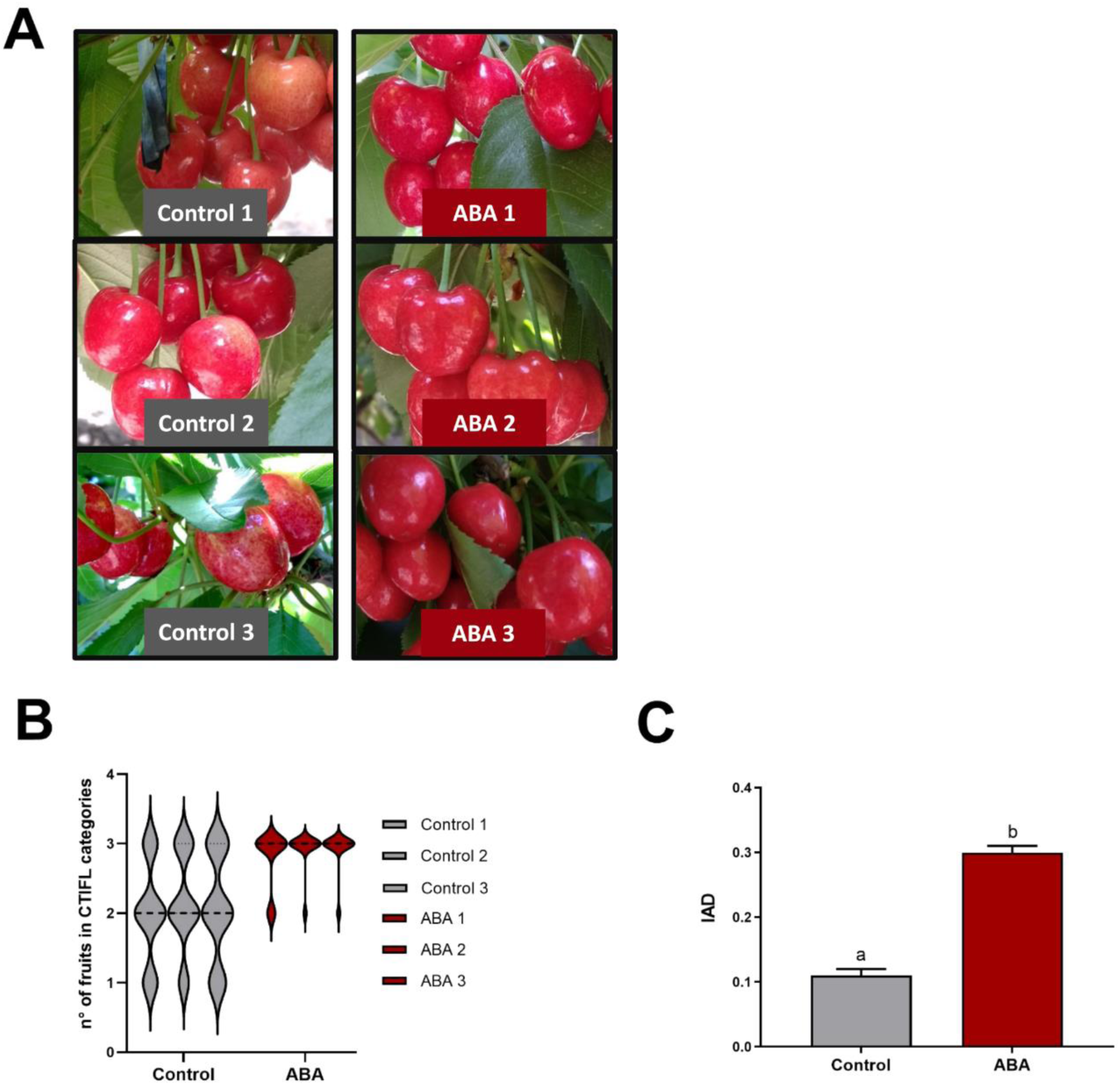
Evaluation of color and ripening parameters in response to ABA treatment. **(A)** Visual color of sweet cherry fruits at 11 days after treatment (DAT) with ABA (67 DAFB), **(B)** Color distribution in different replicates (trees) using CTIFL color chart measured on control and ABA-treated fruits at harvest (13 DAT; 69 DAFB), and **(C)** IAD (index of absorbance difference) at harvest. Three trees as biological replicates were used (20 fruits per tree). In (C), statistical differences were evaluated using a Fisher’s LSD test, where different letters indicate a *p*-value < 0.05. Data are shown as the mean ± SEM.

**Figure 2.**
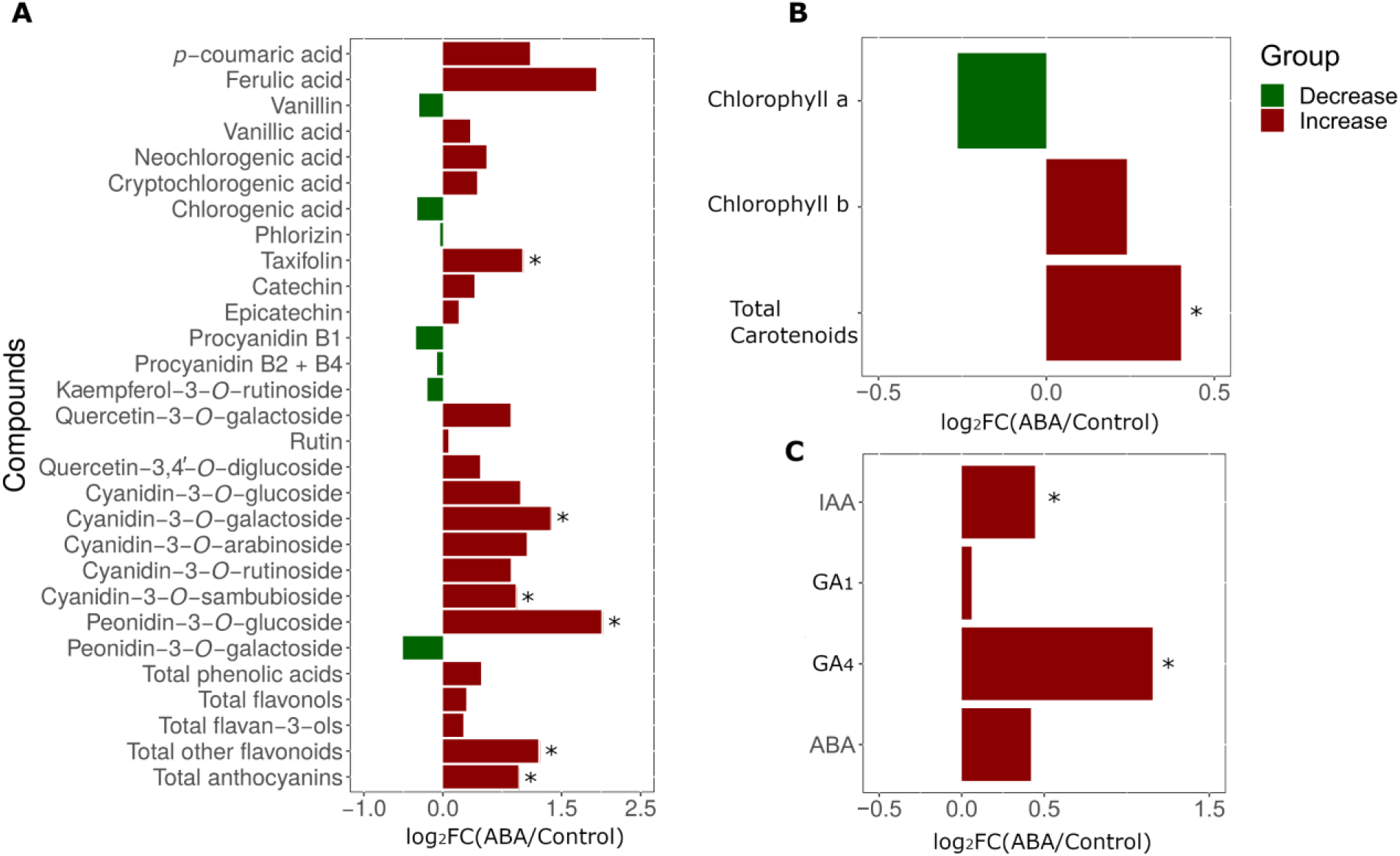
Changes in the accumulation of phenolic and pigment-containing compounds and phytohormones in response to ABA. **(A)** Phenolic compounds, **(B)** Pigment-containing compounds, and **(C)** Phytohormones accumulations were measured at 67 DAFB (11 DAT) in control and ABA-treated fruits. Three trees as biological replicates were used (a pool of at least eight fruits per tree). The data value of each compound was represented as fold change (FC) of the compound concentration between the mean of ABA treated and control fruits in logarithm with base 2 (log_2_FC). The total content of the different flavonoid groups was calculated as the sum of the compounds identified. The other flavonoid groups correspond to taxifolin (5,7,3’,4’-flavan-on-ol) plus phlorizin (dihydrochalcone). Differences were evaluated using a Fisher’s LSD test, where asterisks (*) indicate significant differences at a *p*-value < 0.05 between ABA-treated and control fruits for each compound.

### Genetic and epigenetic modifications during sweet cherry fruit ripening

To understand the modulation of fruit ripening by ABA, we explored the changes occurring during this process at the genetic and epigenetic levels through RNA sequencing (RNA-seq) and Whole Genome Bisulfite Analysis (WGBS), respectively. Sweet cherry fruits at the Early Pink stage (56 DAFB) and at the Red stage (67 DAFB) were investigated with both RNA-seq and WGBS analyses, and fruits from the same trees were used in both analyses. All the replicates (trees) for a given date are grouped in a PCA analysis (data not shown).

RNA-seq of six fruit samples (pools of Early Pink stage fruits from three trees and pools of Red stage fruits from the same three trees) yielded 139 million filtered reads (14.6 Gb of data), with an average length of 100 bp. A total of 123 million reads (88.5% of the filtered read count) were mapped to the P. avium transcriptome reference genome (Supplementary Table S1). Differentially expressed genes (DEGs) were those genes that changed their expression between the Early Pink stage (56 DAFB) and the Red stage (67 DAFB), with a cutoff value of two-fold change (FC) and p-adjusted-value < 0.05. Gene ontology (GO) enrichment analysis of the DEGs is summarized as a Treemap (Fig. 3A, Supplementary Table S2). The GO Biological Process Function head categories that grouped the most genes (p-value < 0.05) were: ‘Response to Abscisic Acid’, ‘Cinnamic Acid Biosynthetic Process’, ‘Sucrose Metabolic Process’, ‘Response to Heat’, ‘Photosynthesis Light Reaction’ and ‘Cell Wall Polysaccharide Metabolic Process’, ‘Cellular Biogenic Amine Catabolic Process’, ‘L-Phenylalanine Metabolic Process’, and ‘Regulation of Dephosphorylation’ (Fig. 3A). Each of these head categories grouped other related subcategories. Additionally, a heatmap of the DEGs from the six most overrepresented GO categories is presented, including upregulated (red) and downregulated (blue) genes (Fig. 3B).

**Figure 3.**
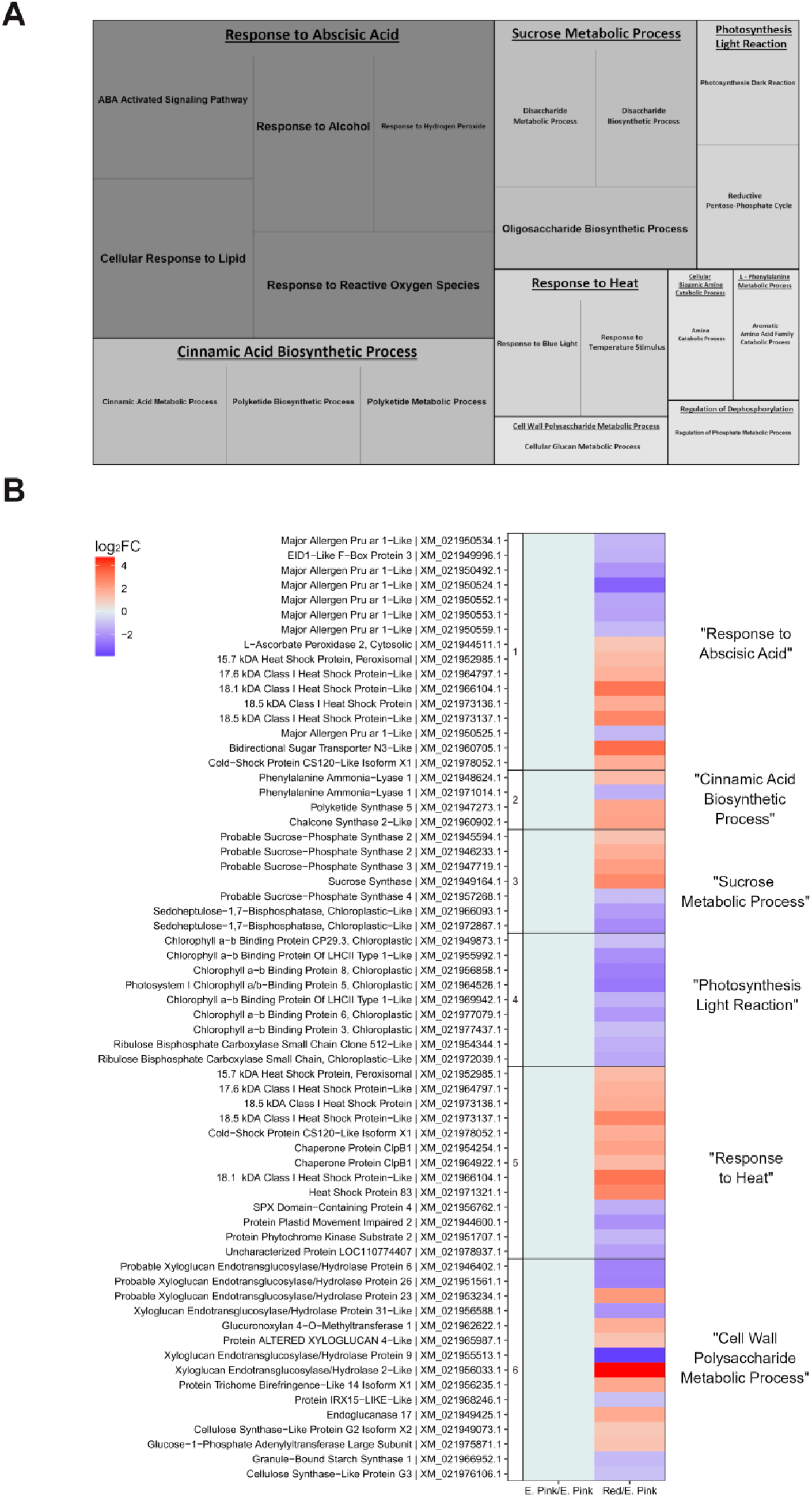
Gene ontology (GO) enrichment analysis of DEGs between two stages of sweet cherry fruit ripening and expression profile of the genes under these categories. **(A)** Treemap representation of overrepresented GO categories among the DEG, and **(B)** heatmap expression profile of the genes from the six most overrepresented GO categories (ReviGO). The biological process function ontologies with the lowest *p*-value and FDR less than 0.01 are shown. For the heatmap, the data value of each compound was represented as fold change (FC) between Red stage samples (67 DAFB) and Early pink stage samples (56 DAFB) in logarithm with base 2 (log_2_FC). Red indicates up-regulation, and blue indicates down-regulation between the comparison stages. All genes are differentially expressed (*p*-adjusted-value < 0.05 and FC > 2). The Treemap and heatmap were designed in R.

WGBS (performed using the same samples as the RNA-seq analyses) yielded a total of 229 million filtered reads (15.7 Gb of data), with an average length of 150 bp and 114 million mapped reads to the P. avium transcriptome reference (49.8% of the filtered read count) (Supplementary Table S3). Both the Early Pink and Red stages of fruit development contained the most abundant CHH context, 46.6% and 48.7% of methylated 5’ cytosines, respectively (Fig. 4A). But, there was an increase in total methylation as fruit color intensifies since the total methylation contexts (CpG, CHG, and CHH) was 4% higher in the Red stage fruits when compared to the Early Pink stage (Fig 4B).

**Figure 4.**
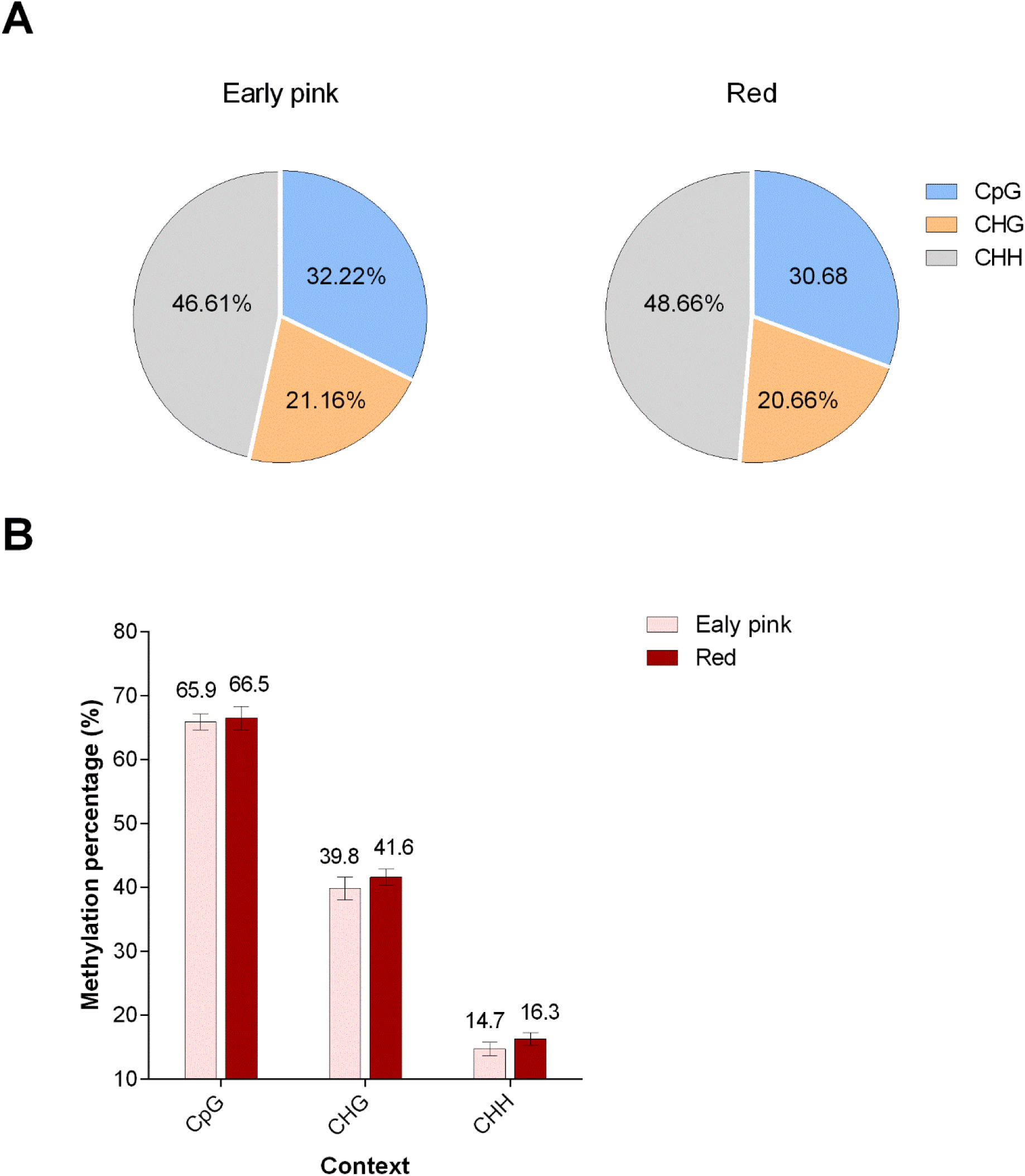
Changes in methylation context during ripening of sweet cherry var. Lapins. **(A)** Changes in the proportion of methylated 5’ cytosines by context (CpG, CHG, CHH) comparing Early pink stage (56 DAFB) and Red stage (67 DAFB), and **(B)** Percentage of 5’ cytosine methylated by context comparing Early pink stage and Red stage.

DMGs (differentially methylated genes) were classified as those genes that presented a DMR (differentially methylated region -percentual change (%) > 15% of DNA methylation in the 5’ cytosines within with q-value < 0.05 between the Early Pink and the Red stage), in the gene or a contiguous region considering the 2,000 bp 5’UTR region upstream the transcription start site. GO analysis was performed on DMGs and represented in a Treemap (Fig. 5A, Supplementary Table S4). The GO Biological Process function categories that grouped the most genes (p-value < 0.05) included the head categories: ‘Cell Wall Polysaccharide Metabolic Process’, ‘Cellular Response to Reactive Oxygen Species’, ‘Carotene Metabolic Process’, ‘Chlorophyll Catabolic Process’, ‘dTDP-Rhamnose Metabolic Process’, ‘Cellular Response to Nutrient’ and ‘COPII-Coated Vesicle Budding’ (Fig. 5A). These head categories grouped other related subcategories. Additionally, a heatmap of the DMGs from the four most overrepresented GO categories is presented (Fig. 5B), including genes hypermethylated (red) and hypomethylated (blue) in the DMRs.

**Figure 5.**
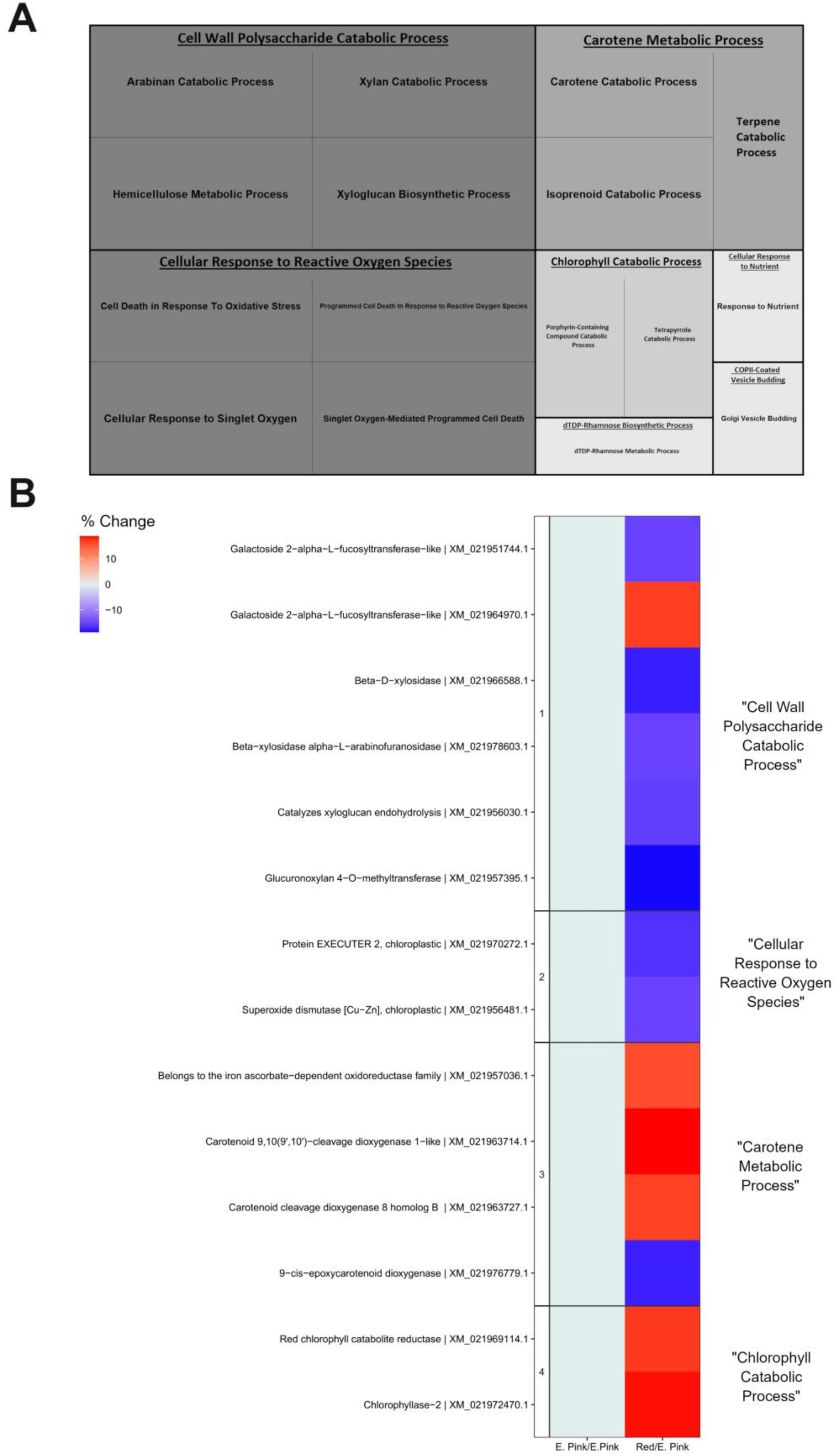
Gene ontology (GO) enrichment analysis performed on DMGs between two stages of sweet cherry fruit ripening and methylation profile of the genes under these categories. **(A)** Treemap representation of overrepresented GO categories, and **(B)** heatmap methylation profile of the genes from the five most overrepresented GO categories (ReviGO). The biological process function ontologies with the lowest *p*-value and FDR less than 0.05 are shown. For the heatmap, the data value of each compound was represented as a percentual change between the Red stage samples and Early pink stage samples. The red color indicates hyper-methylation, and the blue indicates hypo-methylation between the comparison stages. Differentially methylated genes (DMGs) are contiguous (up to 2000 bp) to a differentially methylated region (DMRs) (*q*-value < 0.05 and percentual change > 15%). The Treemap and heatmap were designed in R.

Comparing the DEG and the DMG between the Early Pink and Red stages of fruit development, 46 genes were identified as both differentially expressed and differentially methylated (Fig. 6A). This indicates that 4% of the differentially expressed genes were differentially methylated. Hypermethylated DEGs were downregulated (14 genes) or upregulated (15 genes); similarly, hypomethylated DEGs were downregulated (10 genes) or upregulated (7 genes). Most hypomethylated and hypermethylated genes in their DMRs (23 genes) were not differentially expressed (Fig. 6A, Supplementary Table S5). Only one hypomethylated and hypermethylated gene was upregulated, coding for a TMV resistance protein N-like gene (XM_021977537.1).

**Figure 6.**
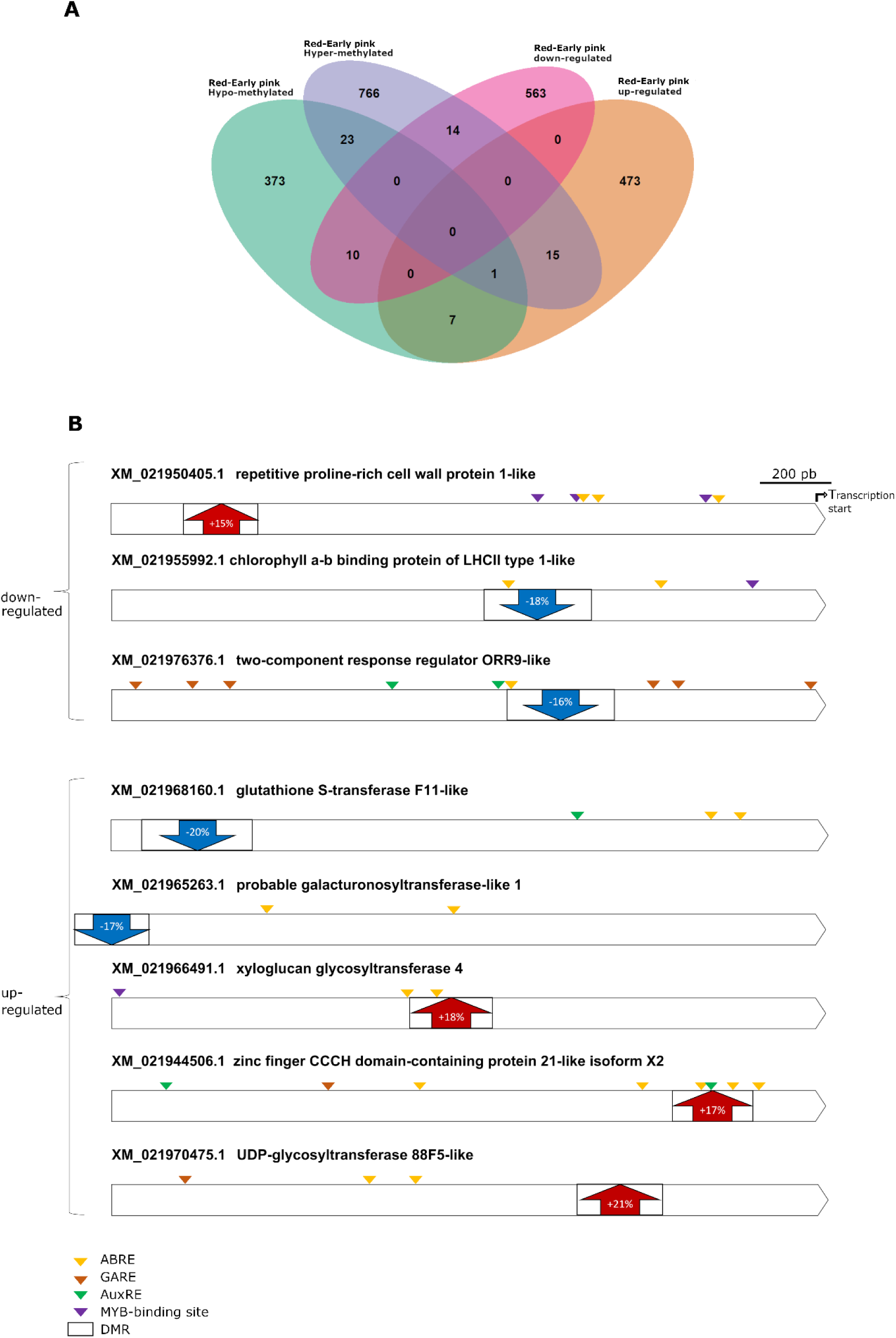
Genes differentially methylated and expressed during ripening with potential hormone-responsive cis-elements in their promoter region. **(A)** Venn diagram between WGBS and RNA-seq analyses during ripening (comparison between Red stage and Early pink stage), and **(B)** Promoter region of selected genes that are differentially expressed and methylated during ripening. Differential expression and methylation according to Table S5. The promoter region was defined as the -2,000 bp from the start of transcription and retrieved from NCBI database. Within the DMR, a black arrow represents hypomethylation (downwards) or hypermethylation (upwards). The percentual change was defined in Fig. 5. Hormone-responsive *cis*-elements were predicted using the PlantCARE database. The Venn diagram was designed in R. The representation of promoter regions was designed using exon-intron GraphicMaker and modified using Gimp v2.10 software.

The DMRs within the -2,000 bp from the start site of representative genes are presented, including their percentage change in methylation in the Early Pink-Red stage comparison; also, some hormone cis-responsive elements within this region are shown (Fig. 6B, Supplementary Tables S5-S6). Chlorophyll a-b binding protein of LHCII type 1-like and the two-component response regulator ORR9-like are downregulated genes in the Early Pink-Red stage comparison having DMRs that decrease their DNA methylation of 5’ cytosines (18% and 16% reduced methylation, respectively) when fruits transition to the Red stage. In contrast, other genes were upregulated with hypermethylated DMRs in the Early Pink-Red comparison: xyloglucan glycosyltransferase 4, zinc finger CCCH domain-containing protein 21-like isoform X2, and the UDP-glycosyltransferase 88F5-like genes (18%, 17%, and 21%, respectively).

### Genetic and epigenetic changes upon ABA treatment at the ripening initiation

Considering that ABA modulates physiological features of ripening (Figs. 1 and 2) and that some genes that change during ripening are potentially ABA-responsive (Fig. 6B, Supplementary Table S6), we performed an RNA-seq and WGBS analyses of fruit samples treated with ABA at the Early Pink stage (56 DAFB). Fruits from trees at the Early Pink stage before the ABA treatment (56 DAFB, T0) and at the Red stage (11 days after the treatment, 67 DAFB, T11) were analyzed through RNA-seq and WGBS analyses. Fruits from the same trees were used in both analyses. In the RNA-seq analyses with ABA treatment, two additional time points were included: 2 h after the treatment (T2h) and 4 d after the treatment (T4).

RNA-seq of the ABA treatment (24 samples, 12 control and 12 ABA-treated from Early Pink and Red stages) yielded a total of 588 million filtered reads (61.4 Gb of data), with an average length of 100 bp and 543 million mapped reads to the P. avium transcriptome reference, which was 92.4% of the total filtered read count (Supplementary Table S7). In this RNA-seq analysis, differentially expressed genes (DEGs) were those genes that changed their expression between T0 (56 DAFB) and T11 (67 DAFB) in every pairwise comparison, with a cutoff value of two-fold change (FC) and p-adjusted-value < 0.01.

When CT11-CT0 DEGs were contrasted with ABAT11-ABAT0 DEGs, some genes were differentially expressed specifically in the ABA-treated samples in the Early Pink to Red transition (175 upregulated and 155 downregulated genes). In contrast, others were differentially expressed only when ABA was not applied (547 upregulated and 404 downregulated genes). In summary, these results revealed a total of 1,281 differentially expressed genes in response to the ABA treatment (ABA-regulated genes) (Fig. 7). The genes in the intersections (320, 1, 1 and 431) changed in both control and ABA trees; these 753 genes appear to be unresponsive to ABA treatment. Categories associated with these T11-T0 ABA-regulated DEGs were considered for GO analysis shown in Fig. 8 (Supplementary Table S8). The Treemap shows that the biological process function ontologies that grouped the most included the head categories ‘Response to Hormone’, ‘Amino Sugar Catabolic Process’, ‘Malate transmembrane transport’, ‘Phosphate Ion Transmembrane Transport’, ‘Nuclear DNA Replication’, and ‘Response to External Biotic Stimulus’. These head categories grouped other related subcategories (Fig. 8A). The five most overrepresented GO categories are presented in the heatmap of the DEGs, including genes that were upregulated (red) and downregulated (blue) (Fig. 8B).

**Figure 7.**
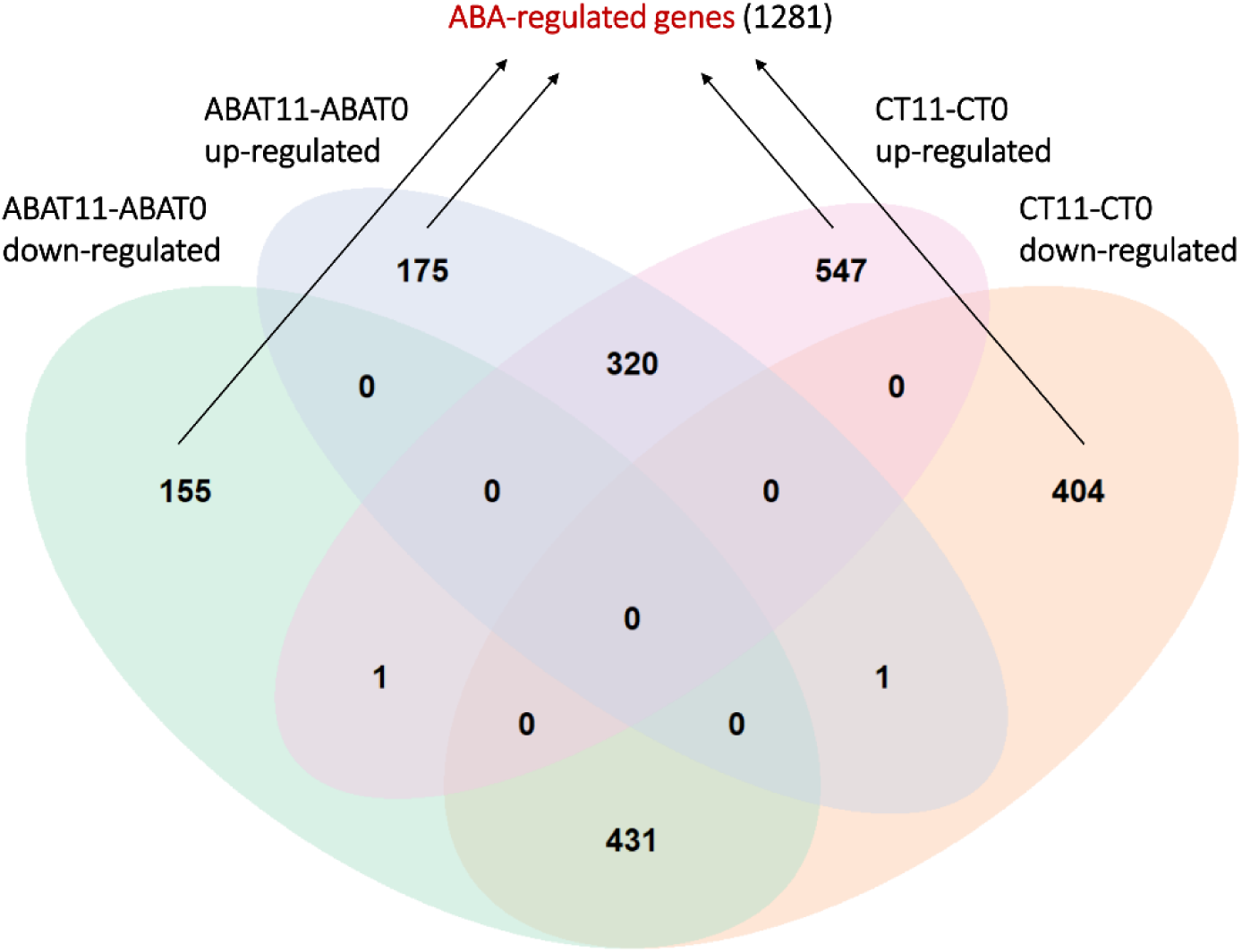
ABA-mediated changes in gene expression in the Red/Early pink comparison (T11/T0). Venn diagram showing the number of DEGs, up- or down-regulated, identified in the pairwise comparisons ABAT11-ABAT0 (ABA-treated) and CT11-CT0 (Control trees). T0 (Early pink stage) samples were collected minutes before ABA treatment, and T11 (Red stage) samples were collected 11 days after the treatment. DEGs were defined as having an absolute fold change value of at least 2.0 and a *p*-adjusted-value less than < 0.05. Venn diagram was designed in R.

**Figure 8.**
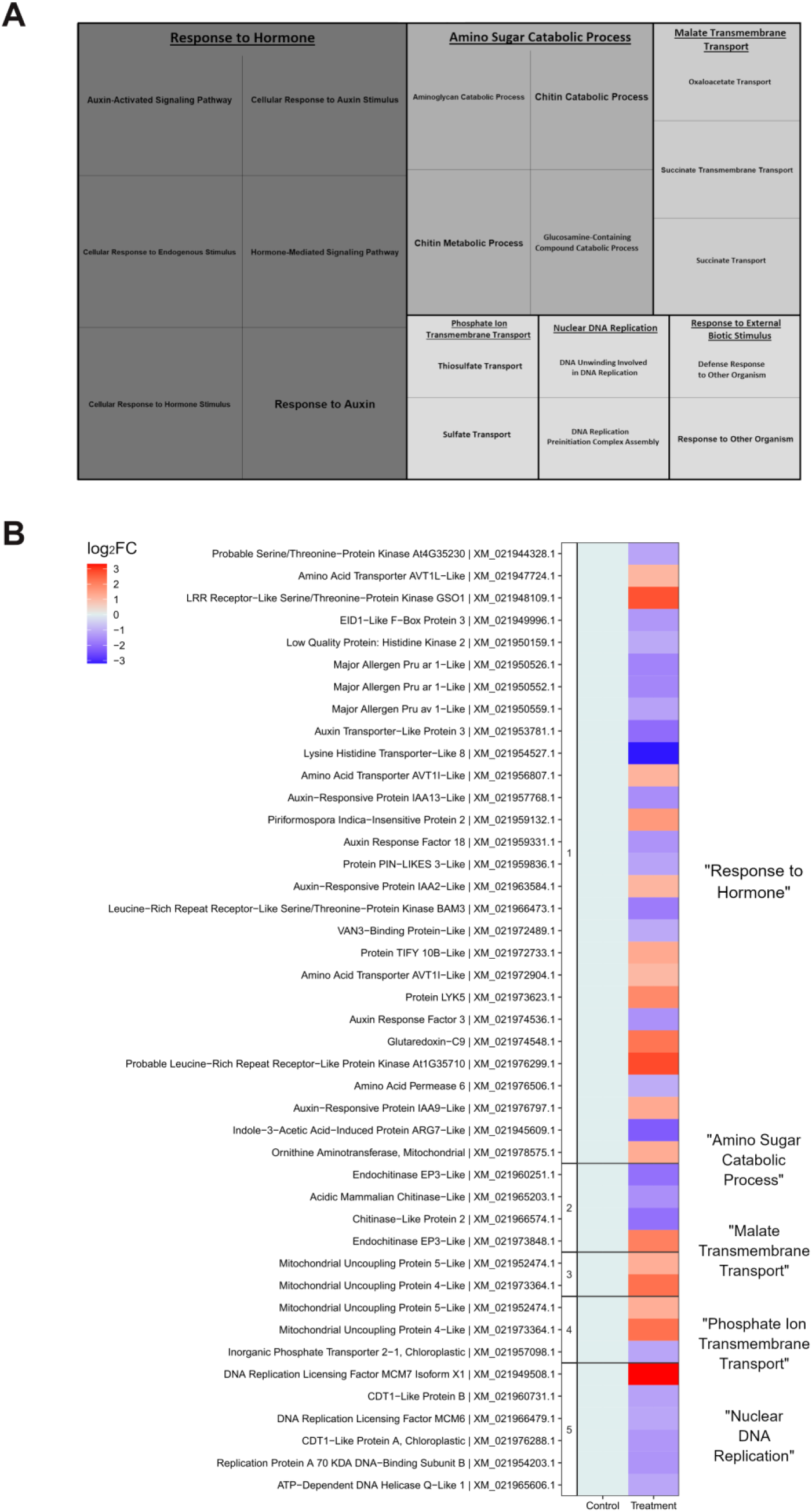
Gene ontology (GO) enrichment analysis was performed on ABA-regulated genes of the Red/Early pink comparison and expression profile of the genes under these categories. **(A)** Treemap representation of overrepresented GO categories, and **(B)** heatmap expression profile of the genes from the five most overrepresented GO categories (ReviGO). The biological process function ontologies with the lowest *p*-value and FDR less than 0.01 are shown. For the heatmap, the data value of each compound was represented as fold change (FC) between Red stage samples (67 DAFB) and Early pink stage samples (56 DAFB) in logarithm with base 2 (log_2_FC). The red indicates up-regulation, and the blue indicates down-regulation, between the comparison stages. All genes are differentially expressed (DEGs) (*p*-adjusted-value < 0.05 and FC > 2.0). The Treemap and heatmap were designed in R.

A similar analysis was performed in the T4-T0 comparison, showing some GO categories in common with T11, including ‘Response to Hormone’, and ‘Amino Sugar Catabolic Process’ (Supplementary Figure S2, Supplementary Table S9). Additionally, the T2h-T0 comparison had overrepresented the molecular function GO category “DNA Binding” that showed changes in genes coding for MYB, WRKY, NAC, ERF transcription factors, and changes in gene models encoding histone proteins (Supplementary Figure S3).

WGBS of the ABA treatment (six samples, three control and three ABA-treated at the Red stage) yielded a total of 226 million filtered reads (15.5 Gb of data), with an average length of 150 bp and 113 million mapped reads to the P. avium transcriptome reference, which was 50.3% of the total filtered read count (Supplementary Table S10). GO analysis was performed on DMGs, represented in a Treemap, after reducing and grouping in semantic similarities of the categories (Fig. 9A, Supplementary Table S11). The biological process function ontologies that grouped the most included the head categories ‘Neutral Lipid Biosynthetic Process’, ‘Cellular Hormone Metabolic Process’ and ‘Chlorophyll Catabolic Process’ (Fig. 9A). A heatmap of the DMGs from the four most overrepresented GO categories is showed (Fig. 9B), including genes hypermethylated (red) or hypomethylated (blue) in the DMRs.

**Figure 9.**
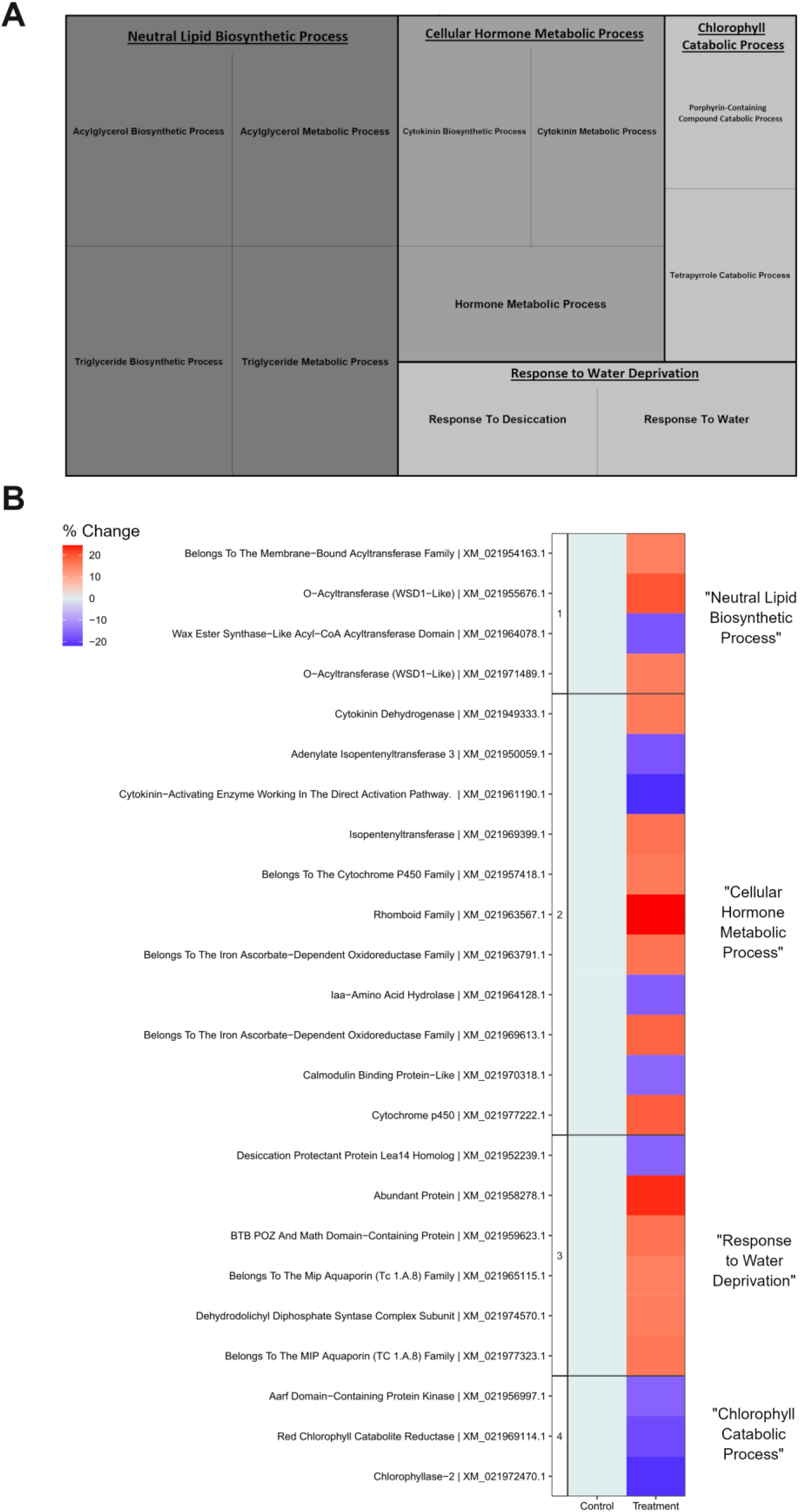
Gene ontology (GO) enrichment analysis was performed on ABA-responsive DMGs in the Red stage and the methylation profile of the genes under these categories. **(A)** Treemap representation of overrepresented GO categories, and **(B)** heatmap methylation profile of the genes from the five most overrepresented GO categories (ReviGO). The biological process function ontologies with the lowest *p*-value and FDR less than 0.05 are shown. For the heatmap, the data value of each compound was represented as a percentual change between ABA-treated fruits and Control fruits at the Red stage (ABAT11-CT11). The red color indicates hyper-methylation, and the blue indicates hypo-methylation between the comparison stages. Differentially methylated genes (DMGs) are contiguous (up to 2000 bp) to a differentially methylated region (DMRs) (*q*-value < 0.05 and percentage change > 15%). The Treemap and heatmap were designed in R.

## Discussion

### ABA modulates physiological changes during ripening in sweet cherry

ABA triggers ripening in non-climacteric fruit species (Fuentes et al., 2019; Setha, 2012). Its effect occurs on almost all ripening-related parameters, with anthocyanin accumulation in the fruit peel and related color acquisition being the most evident ABA-driven events, as reported in colored fruits, such as strawberry, grapevine, and sweet cherry (Jiang et al., 2003; Jia et al., 2011; Ren et al., 2011; Shen et al., 2014; Sun et al., 2010; Gagné et al., 2011). Here we show increased color accumulation (Fig. 1A) and a more homogeneous color distribution (Fig. 1B), suggesting that ABA advances the coloring process. ABA influence or ripening-related parameters was previously reported in grapevine, where monitoring over color and °Brix parameters was performed during the ripening process (Wheeler et al., 2009). In sweet cherry we recently reported that ABA advances the increase in IAD - a ripening index - in sweet cherry fruits (Kuhn et al., 2021a). Here, we report higher IAD at harvest, similar to other cultivars (Time et al., 2021). Regarding the other ripening parameters, we found reduced size, weight, firmness, and sugar content (Supplementary Fig. S1). Previous works showed that exogenous ABA promoted sugar accumulation (Ren et al., 2011; Wang et al., 2015), which differs from our results. Possibly, this is associated with the ABA application, as in our case, whole trees were treated with ABA in planta, whereas the previously mentioned studies were performed ex planta (excised shoots bearing fruits or excised fruits treated with an ABA solution), where the natural source-sink interactions are altered. Luo et al. (2014) showed that in planta, ABA-treated fruits had a larger size and increased soluble solids content, but it is unclear if the leaves received the ABA treatment. In this regard, Time et al. (2021) showed that fruits treated in planta and dipped in ABA solution (thus, excluding the leaves) had fewer soluble solids than untreated samples. Therefore, the effect of ABA on fruit sugars seems highly dependent on its influence on leaves and possibly in the source-sink interactions. Regarding softening, fruit firmness is reduced during ripening and in response to ABA (Luo et al., 2014; Ren et al., 2011; Time et al., 2021), which is in line with our results.

Regarding marker molecules, exogenous ABA increases the fruit’s endogenous ABA and anthocyanin content (Luo et al., 2014; Kuhn et al., 2021a; Ren et al., 2010; Shen et al., 2014; Sun et al., 2010; Wheeler et al., 2009). In our experiment, the ABA treatment increased the ABA content in the fruits 11 days after the treatment and increased the content of several anthocyanins considerably(Figs. 2A and 2C). Other plant hormones, such as IAA and gibberellins, are antagonistic hormones of ripening (Böttcher et al., 2010; Teribia et al., 2016; Kuhn et al., 2020; Kuhn et al., 2021b). ABA reduces the IAA content when applied before color initiation in sweet cherry (Wang et al., 2015). Our results differ from these previous findings, as we found that ABA increases IAA content (Fig. 2C). This is possibly due to the timing of the treatment, as we applied ABA to the trees when the fruits were already colored (10%-15% of the fruits in the trees were pink colored) and possibly at this stage the ABA-dependent IAA increase might prevent an excessive ABA effect.

Carotenoids accumulate during ripening in several fruit species (Beekwilder et al., 2008; Karppinen et al., 2016; Leng et al., 2017). We found that ABA increased the content of carotenoids (Fig. 2B), which is in line with previous reports on tomatoes (Barickman et al., 2014; Barickman et al., 2016).

As the fruits ripen, the green color disappears, and the photosynthetic rate decline. In this regard, chlorophyll is a negative molecular marker. For instance, during grapevine fruit ripening, chlorophyll a and b are reduced (Leng et al., 2017); similarly, total chlorophyll content declines during sweet cherry fruit development (Shen et al., 2014). We found that ABA modulates both chlorophyll types (Fig 2B). ABA promotes chlorophyll degradation during leaf senescence (Gao et al., 2016); thus, reduced chlorophyll a is expected.

### Fruit ripening involves changes at the genetic and epigenetic level that underlies physiological modifications

In the Pink to Red stage transition, several overrepresented categories in the GO analysis of the RNA-seq reflected the physiological changes reported during ripening, including the ‘Cinnamic Acid Biosynthesis Process’, ‘Sucrose Metabolic Process’, ‘Photosynthesis Light Reaction’ and ‘Cell Wall Polysaccharide Metabolic Process’ categories (Fig. 3A). The genes of the ‘Cinnamic Acid Biosynthesis Process’ category include the phenylalanine ammonia-lyase 1 and the chalcone synthase 2-like gene models (Fig 3B). PacCHS expresses during the coloring process of sweet cherry, differentially expressing between dark and pink cherries (Liu et al., 2013). Moreover, PacCHS, an essential gene for the coloring process that occurs during ripening, is significantly correlated with anthocyanin accumulation in dark varieties (Liu et al., 2013). Additionally, we found upregulation and hypermethylation of UDP-glycosyltransferase 88F5-like (KEGG entry 100853970) (Fig. 6B), an ortholog of PGT (Phlorizin synthase, that synthesizes a dihydrochalcone; Judgé et al. 2008), suggesting that this pathway is also regulated at an epigenetic level.

Regarding the ‘Sucrose Metabolic Process’, sucrose phosphatase synthase and sucrose synthase gene models were upregulated in Red stage fruits compared with Pink stage fruits (Fig. 3B). This agrees with the well-reported increase in sugar content in non-climacteric fruits (Kuhn et al., 2014; Luo et al., 2014). Regarding the ‘Photosynthesis Light Reaction’ category, several genes encoding chlorophyll-binding proteins and two genes coding for a ribulose bisphosphate carboxylase small chain were downregulated in the Pink to Red stage transition (Fig. 3B); this agrees with the reduction in chlorophyll content during the progression of fruit ripening (Shen et al., 2014). Finally, in the ‘Cell Wall Polysaccharide Metabolic Process’ category, several gene models encoding xyloglucan endotransglucosylase/hydrolase (XTH) had differential expression (Fig. 3B). XTHs are key cell wall remodeling enzymes, and their encoding genes change during fruit development (Opazo et al., 2017). For instance, in tomato fruits, important xyloglucan depolymerization occurs during softening, accompanied by an increase in the expression of SlXTH5 and SlXTH8 (Miedes et al., 2009). In strawberries, higher expression of FvXTH9 and FvXTH6 enhances fruit ripening (Witasari et al., 2019).

In the ‘Response to Heat’ category, several genes encoding heat shock-related proteins, including some chaperone proteins, were identified (Fig. 3B). In tomatoes, several genes encoding Heat Shock Proteins (HSPs) superfamily proteins express differentially during ripening (Bertero et al., 2021). Their expression is specific to different ripening processes; for instance, HSP70 is involved in carotenoid synthesis in tomato fruits (D’Andrea and Rodriguez-Concepcion, 2019).

ABA biosynthesis and catabolism are important during the ripening process of non-climacteric fruits (Liao et al., 2018). ABA response includes several pathways related to development and stress responses. We found several genes grouped in the ‘Response to Abscisic Acid’ category, including some that encode for HSPs (Fig. 3B). Remarkably, a gene encoding a bidirectional sugar transporter N3-like was highly upregulated in this category.

Ripening is a process that involves changes in the DNA epigenetic status. For instance, in the non-climacteric sweet pepper and strawberry, DNA hypomethylation occurs in the upstream region of the transcriptional start site of several ripening-related genes during fruit ripening (Xiao et al., 2020; Cheng et al., 2018). In contrast, in orange, DNA hypermethylation is required for this process to occur (Huang et al., 2019). In agreement with the work on orange, we found that global DNA methylation increased during ripening, as Red stage fruits had higher methylation % (Fig. 4B).

The bisulfite sequencing revealed several categories overrepresented in the GO analysis of genes differentially methylated in their -2000 bp 5’UTR region between the Pink and Red stages (Fig. 5A). The category ‘Cellular Response to Reactive Oxygen Species’ included a gene encoding a superoxide dismutase (Fig. 5B), an enzyme involved in the dismutation of superoxide radicals (O_2_^−^) to molecular oxygen (O_2_). Reactive oxygen species (ROS) have relevance during ripening. Recently, it was reported that the treatment with H_2_O_2_ advances ripening in grapevine fruits (Guo et al., 2020). Considering changes in ROS, several enzymes that alleviate their levels are expected to increase. In our results, the hypomethylation of the DMRs of the gene encoding this superoxide dismutase might produce changes in its expression, which might influence the defense against oxidative stress.

Additionally, the category ‘Carotene Metabolic Process’ included several hypermethylated genes coding for putative carotenoid cleavage dioxygenase (CCD; Fig. 5B). CCDs are involved in white color formation (Ohmiya et al., 2006); therefore, their regulation at the epigenetic level might explain color changes attributed to carotenoids instead of anthocyanins in fruits. A gene model encoding a putative 9-cis-epoxy-carotenoid dioxygenase had a hypomethylated DMR (Fig. 5B). Upregulation of PavNCED1 gene during sweet cherry ripening was reported (Kuhn et al., 2021a; Ren et al., 2011).

Besides the transcript variation of genes involved in the xyloglucan metabolism during ripening (Fig. 3), some genes, including xylosidases, had DMRs differentially methylated (Fig. 5B), suggesting that the cell wall loosening is a process controlled genetically and epigenetically. The “xyloglucan glycosyltransferase 4 encoding gene” expression has an upregulation in gene expression and a hypermethylated DMR in the Pink-Red stage comparison (Fig. 6B). This gene also contains ABA-responsive elements (Fig. 6B).

Other genes with differential expression and methylation are the chlorophyll a-b binding protein of LHCII type 1-like and the two-component response regulator ORR9-like, which is related to the cytokinin response pathway (Fig. 6B).

### ABA modulates pathways related to fruit ripening at the genetic and epigenetic level

Our results show that the progression of ripening is strongly correlated to ABA at the physiological and genomic levels. Some ABA response genes activate during ripening (Fig. 3B), and some genes that change during ripening have ABA-responsive elements (Fig. 6B); thus, this hormone is an integral part of the ripening process. Therefore, we studied the transcriptomic profile and DNA methylation pattern of ABA-treated fruits to understand how this hormone modulates ripening in sweet cherry fruits.

RNA-seq analysis was performed in fruits treated with ABA and control fruits within the temporal window comprised between color initiation (Pink stage; T0) and red color (Red stage, T11). A GO functional analysis examined the ABA-regulated genes defined in Fig. 7. Therefore, the genes associated with these GO terms were subsequently analyzed (Fig. 8). We found some ABA-regulated genes encoding chitinases and endo-chitinases with differential expression between T0 and T11 (Fig 8B). Chitinases are well-described as ripening markers. For instance, in grapevine, the expression of chitinase genes is strongly upregulated during ripening, coinciding with sugar accumulation (Robinson et al., 1997). Chitin is part of the cell wall; therefore, an upregulation of chitinase expression during fruit softening is expected.

Hormones are essential in the control of ripening. We found several differentially expressed genes associated with the ‘Response to Hormone’ category encoding putative auxin-responsive proteins from the Aux/IAA family and auxin response factors (Fig. 8B). The involvement of the most abundant auxin, IAA, in the fruit ripening process as antagonizing hormone is well described in grapevine (Böttcher et al., 2010; Dal Santo et al., 2020). In addition, an ABA-IAA switch has been proposed in this species to control the ripening initiation time (Gouthu and Deluc, 2015). In sweet cherry, IAA affects different genes from the ABA pathway at the color initiation stage (Wang et al., 2015). Therefore, a link between ABA, auxin, and the ripening initiation time seems to exist. Our results suggest a possible role of ABA in ripening modulation by controlling the expression of the auxin response pathway. This pathway is also influenced by gibberellin in sweet cherry (Kuhn et al., 2021b), so a possibility arises where ABA and gibberellin control ripening initiation by regulating auxin response genes.

Further studies are required to decipher this regulatory module. Genes encoding an auxin transporter-like protein 3 and a protein PIN-LIKES 3-like were also downregulated (Fig 8B). Auxin transport might have some relevance during fruit ripening, and it is proposed that less auxin transport from the fruits increases berry coloring (Serrano et al., 2022). Therefore, we suggest that ABA’s control of auxin transport-related genes might be relevant during ripening initiation.

Concerning the hormonal control of ripening at the epigenetic level, we identified several genes associated with the cytokinin biosynthesis process differentially methylated in their DMRs in response to ABA (control fruits vs. ABA-treated fruits at T11). These genes encode isopentenyl transferases and cytokinin dehydrogenase (Fig. 9B). There are few studies about the role of cytokinins in non-climacteric fruit ripening, although they have been shown to play a role in climacteric fruit ripening (Mujica et al., 2020). In grapevine, the content of the cytokinin isopentenyladenine (iP) increases during fruit ripening (Böttcher et al., 2015), whereas in sweet cherry, both 2-iP and isopentenyladenosine increase as the fruits turn red (Teribia et al., 2016). In ‘Shine Muscat’ grapevine fruits, chlorophyll better endured the treatment with CPPU (N-(2-chloro-4-pyridyl)-3-phenylurea) at veraison (Suehiro et al., 2019). In grapevine, CPPU applied to small green fruits reduced color development, anthocyanin content, and sugar accumulation (Tyagi et al., 2021; Tyagi et al., 2022). These works and our results suggest that cytokinin plays a role during non-climacteric ripening, modulated by ABA at the epigenetic level, possibly acting as a ripening antagonizing hormone. In this regard, we also found a two-component response regulator ORR9-like that is upregulated and hypermethylated during ripening, with several hormone-responsive cis-elements in their promoter region, showing that not only the biosynthesis pathway may be regulated at an epigenetic level but also the response pathway of cytokinin.

Chlorophyll catabolism is a characteristic of fruit ripening. Chlorophyll breakdown into cleavage products occurs in consecutive steps catalyzed by chlorophyllases and other enzymes (Goldschmidt et al., 2000). In our results, the ‘Chlorophyll Catabolic Process’ GO term includes differentially methylated genes in their DMRs in the Early Pink to Red stage transition and after the ABA treatment. The gene model chlorophyllase-2, which is ABA-responsive regarding their DNA methylation status (hypomethylated in ABA-treated samples; Fig. 9B), also is hypermethylated during ripening (Fig. 5B). Chlorophyllase is an important enzyme in chlorophyll catabolism (Harpaz-Saad et al., 2007). Thus its epigenetic regulation during ripening and modulation by ABA might contribute to its expression variations and function. These findings align with the observed effect of ABA on reducing the content of chlorophyll a (Fig. 2B).

Fruit ripening is an oxidative phenomenon (Jiménez et al., 2002). Regarding the reactive oxidative species (ROS), changes in H_2_O_2_ and lipid peroxidation occur during the initiation of the red coloration in tomato fruits (Jiménez et al., 2002). ROS marker enzyme activity in this organism, including cytochrome c oxidase and catalase, was reported during fruit ripening (López-Vidal et al., 2016). Under these oxidative conditions, antioxidant systems alleviate the ROS levels, including the glutathione-ascorbate cycle. For instance, glutathione reductase activity increased during tomato fruit ripening (López-Vidal et al., 2016). Glutaredoxins regulate the cellular redox state by using the reducing capacity of glutathione. We found an ABA-regulated gene encoding a putative glutaredoxin-C9 (Fig. 8b), which was upregulated at the Red stage, suggesting that in sweet cherry, the ascorbate– glutathione cycle might play a role during fruit ripening. Similar to was has been seen in grapevine (Guo et al., 2020), two predicted sweet cherry iron ascorbate-dependent oxidoreductase genes had increased DMR methylation in response to ABA treatment (Fig. 9B). This finding suggests that the ascorbate–glutathione cycle might be modulated by ABA at the genetic and epigenetic level during fruit ripening and that ROS may play a role during non-climacteric fruit ripening in sweet cherry.

Ethyl and acetate esters are naturally produced in plants and are responsible for the production of key flavors in fruits (Morton and Macleod, 1990). Wax ester synthases synthesize esters from long-chain acyl-CoA and fatty alcohols. It has been reported that tomato, kiwi, and melon fruits produce these ester synthases (Galaz et al., 2013; Günther et al., 2011; Lin et al., 2016). Bisulfite sequencing demonstrated that the GO category ‘Neutral Lipid Biosynthetic Process’ was overrepresented in ABA-treated samples, and some genes associated with this category were differentially methylated in the 5’UTR region – 2000 bp upstream of the transcription start site (Fig. 9B). This suggests a role for ABA in the development of flavor, a characteristic feature of fruit ripening.

A summary of the findings of this work is included in Fig. 10, comprising overrepresented GO categories modulated at the genetic and epigenetic level during fruit ripening and in response to ABA.

**Figure 10.**
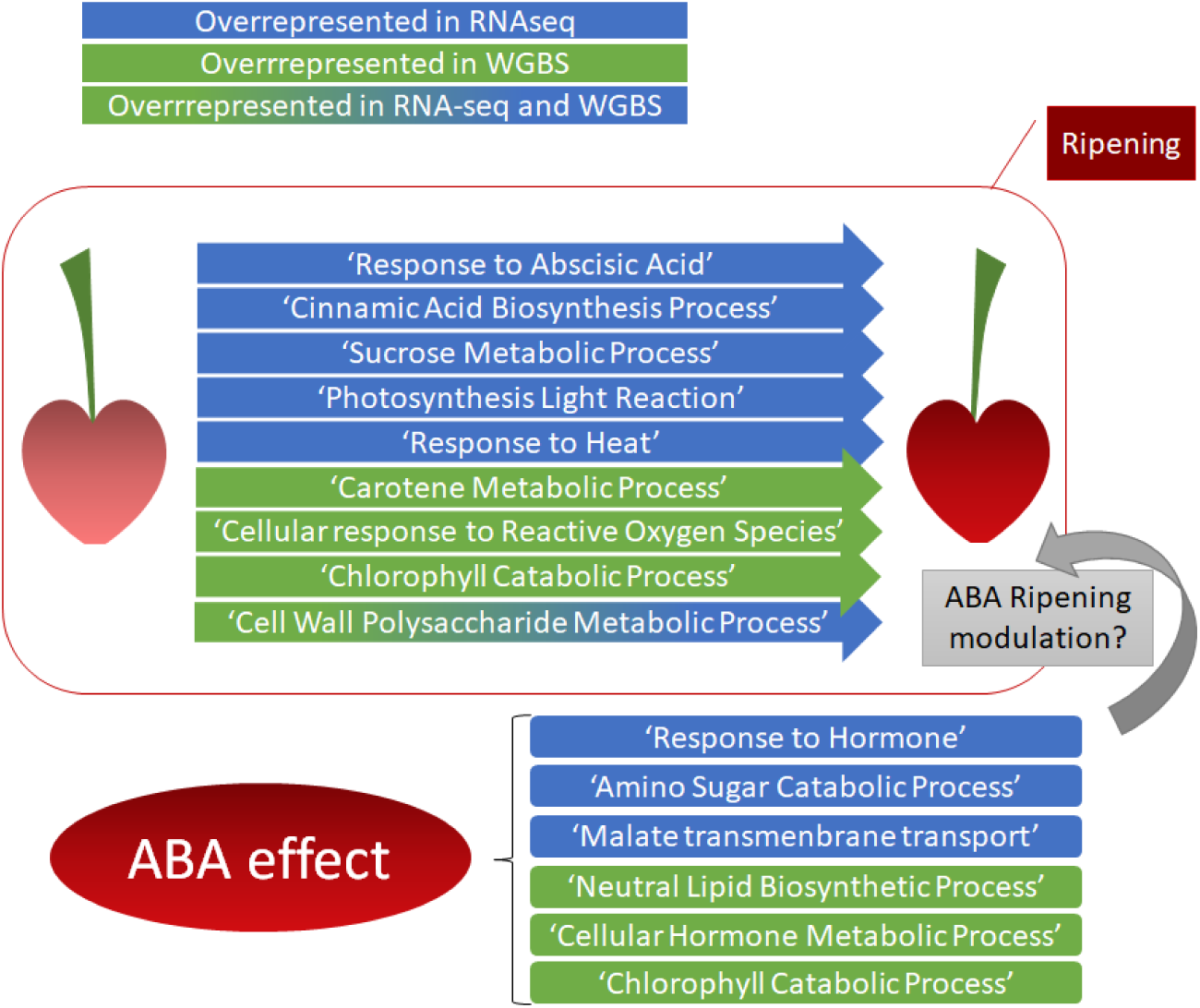
Scheme summarizing the main changes during ripening (Early Pink - Red stage comparison) and in response to ABA treatment regarding the overrepresented categories found in the RNA-seq and WGBS analyses.

## Supplementary data

The following supplementary data are available on JXB online.

**Figure S1.** Evaluation of ripening physiological parameters in response to ABA treatment.

**Figure S2.** Gene ontology (GO) enrichment analysis performed on ABA-regulated genes of the Late Pink/Early pink comparison and expression profile of the genes under these categories.

**Figure S3.** Expression profile of the genes under the overrepresented category “DNA binding”.

**Table S1**. Mapping statistics of ripening RNA-seq analysis

**Table S2.** Summary GO analysis RNA-seq in ripening

**Table S3.** Mapping statistics of ripening WGBS analysis

**Table S4.** Summary GO analysis WGBS in ripening

**Table S5.** Genes that change transcriptionally and in methylation during ripening

**Table S6.** Cis-element identification in methylation-transcriptionally regulated genes during ripening

**Table S7.** Mapping statistics of ABA-treatment RNA-seq

**Table S8.** Summary GO analysis RNA-seq in ABA-treatment T11-T0

**Table S9.** Summary GO analysis RNA-seq in ABA-treatment T4-T0

**Table S10.** Mapping statistics of ABA-treatment WGBS analysis

**Table S11.** Summary GO analysis WGBS in ABA-treatment

**Table S12.** Expression matrix of control plants

**Table S13.** Expression matrix of ABA-treated plants

**Table S14.** WGBS analysis and methylation matrix

## Acknowledgments

We want to thank the Fundo el Parque for access to the commercially produced sweet cherry trees used in these studies. We would also like to thank Alson Time, Simón Miranda, Natalia Molina, and Elisabeth Sarabia for their technical support with the fruit quality measurements.

## Author contributions

NK, MA, CP, and LM performed Conceptualization, Investigation, Methodology and Validation of the research. NK, CP, MA, CH, FC, SM, and EC performed Data curation and Formal Analysis. CP, MA, CH, and FC contributed with Software analyses. NK, CP, MA, and LM performed Visualization of the research. Writing – Original Draft Preparation was performed by NK, MA, CP and LM. Writing – Review and Editing were performed by LM, JMD, and SMA. LM, JM, SMA and NK provided the Resources. LM and NK performed Supervision and Project Administration. LM provided the Funding Acquisition.

## Data Availability

The data have been deposited with links to BioProject accession number PRJNA898489 in the NCBI BioProject database (https://www.ncbi.nlm.nih.gov/bioproject/).

## Abbreviations

ABA: Abscisic acid
DAFB: Days after full bloom
DEGs: Differentially expressed genes
DMGs: Differentially methylated genes
DMR: Differentially methylated region
GO: Gene Ontology
IAD: Index of absorbance difference
RNA-seq: RNA sequencing
UHPLC-MS/MS: Ultraperformance liquid chromatography-tandem mass spectrometry
WGBS: Whole-genome bisulfite sequencing

